# Cell Geometry Distinguishes Migration-Associated Heterogeneity in Two-Dimensional Systems

**DOI:** 10.1101/2021.12.22.473305

**Authors:** Sagar S Varankar, Kishore Hari, Sharmila A Bapat, Mohit Kumar Jolly

## Abstract

**Background:** *In vitro* migration assays are a cornerstone of cell biology and have found extensive utility in research. Over the past decade, several variations of the two-dimensional (2D) migration assay have improved our understanding of this fundamental process. However, the ability of these approaches to capture the functional heterogeneity during migration and their accessibility to inexperienced users has been limited.

**Methods:** We downloaded published time-lapse 2D cell migration datasets and subjected them to feature extraction with the Fiji software. We used the ‘Analyze Particles’ tool to extract ten cell geometry features (CGFs), which were grouped into ‘shape’, ‘size’ and ‘position’ descriptors. Next, we defined the migratory status of cells using the ‘MTrack2’ plugin. All data obtained from Fiji were further subjected to rigorous statistical analysis with R version 4.0.2.

**Results:** We observed consistent associative trends between size and shape descriptors and validated the robustness of our observations across four independent datasets. We used these descriptors to resolve the functional heterogeneity during migration by identifying and characterizing ‘non-migrators (NM)’ and ‘migrators (M)’. Statistical analysis allowed us to identify considerable heterogeneity in the NM subset, that has not been previously reported. Interestingly, differences in 2D-packing appeared to affect CGF trends and heterogeneity of the migratory subsets for the datasets under investigation.

**Conclusion:** We developed an analytical pipeline using open source tools, to identify and morphologically characterize functional migratory subsets from label-free, time-lapse migration data. Our quantitative approach identified a previously unappreciated heterogeneity of non-migratory cells and predicted the influence of 2D-packing on migration.

## Introduction

Migration is a fundamental cellular function which contributes to development, tissue maintenance and disease progression across biological systems. Initiated during early embryonic morphogenesis, migration involves dynamic changes to the cellular architecture and cell-niche interactions^1–3^. Apart from well-defined molecular signatures, phenotypic parameters are often used to identify migrating cells albeit with limited accuracy during homeostasis. Changes in cell phenotype or, from a quantitative perspective, geometry are most evident during processes like the epithelial-mesenchymal transition (EMT) or immune cell homing which involve shifts between distinct cellular states^4^. Dependent grossly on tissue architecture and spatial arrangement, cells can adopt a defined repertoire of geometric configurations which in turn influence their patterns of motility^5–8^. While the physiological relevance of distinct migratory modalities has been widely studied *in vitro*, the functional heterogeneity of migration in the context of cell geometry remains poorly understood.

The *in vitro* two-dimensional (2D) scratch assay is routinely used to quantify and examine the effect of exogenous cues on cell migration^9^. In recent years, several alterations to its experimental setup in conjunction with quantitative tools have provided insight into the biophysical properties of migrating cells^10–15^. In a previous study, we coupled time lapse microscopy with the 2D scratch assay and defined three distinct migratory modes in ovarian cancer cell lines, *viz*., passive collective cell migration (pCCM), active collective cell migration (aCCM) and EMT. We further demonstrated the switching of migration modalities in response to extrinsic cues for these cell lines and observed that cells which undertook aCCM exhibited higher chemoresistance and a greater inherent plasticity^16^. A similar study with breast cancer cell lines associated monolayer migratory patterns with specific mutational signatures, thus providing a clinically relevant readout for metastatic potential^17^. Independent reports have further discerned the leader-follower dynamics, cell polarity and orientation, and directionality of collective cell migration with similar experimental approaches^18–20^. Cell tracking and particle image velocimetry (PIV) approaches have also been adopted in tissue-derived scaffolds^21^ and developing embryos^22^ to dissect niche dependent migratory dynamics. In addition to the use of time lapse microscopy, the recent development of high-throughput analytical tools has allowed efficient handling of large migration datasets^13^. Microfluidic devices in conjunction with machine learning have been used to discern the shape-shifting dynamics of single cells^23^. However, despite their utility, most of these platforms require high-end computing and are often inaccessible to users with limited coding experience.

We developed an open source quantitative approach, utilizing cell geometry features (CGFs), for label-free time lapse microscopy data. Our analysis allowed the identification and characterization of migratory subsets within otherwise homogenous cell systems. Interestingly, CGF trends and functional heterogeneity observed in our study extended across multiple experimental models of cell migration. These results allowed us to speculate the effects of 2D-packing, growth factor treatment and ECM coating on migratory subsets, and emphasized the widespread utility of our pipeline. Our approach to conduct image analysis with Fiji and the availability of well-annotated codes would allow experimental biologists to efficiently integrate this pipeline for analysing cell migration data.

## Materials and Methods

### Imaging Datasets, Processing and Quantification of imaging data

Label-free time-lapse imaging data acquired with phase-contrast microscopy were downloaded from published studies^16,24–26^ and processed with Fiji software (https://imagej.net/Fiji/Downloads) as previously described^12^. Briefly, images from individual datasets were converted into an 8-bit format, adjusted for contrast and threshold, and analysed with the ‘Analyze Particles’ tool and ‘MTrack2’ plugin in Fiji. CGFs were extracted across the time course for each experimental dataset. Migratory and non-migratory cells were identified by comparing and merging cell position co-ordinates from the ‘Analyze Particle’ and ‘MTrack2’ outputs in R version 4.0.2. Codes used for data wrangling have been uploaded to Github and can be found at https://github.com/askhari139/CGF.

### Cumulative Correlation Scores

Cumulative correlation scores were calculated as follows:

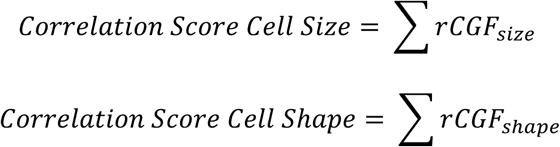

where, rCGFsize = Pearsons’ Correlation coefficient between two Size CGFs

rCGF_shape_ = Pearsons’ Correlation coefficient between two Shape CGFs

Only ‘r’ values with a p<0.05 were used for this calculation

### Statistical analysis

Statistical analyses were performed with Prism software package version 6.0 (GraphPad). *P* values were calculated using two-tailed paired Student’s t test and considered significant if *P*<0.05. Pearson’s Correlation Co-efficient and Principal Component Analysis were calculated in R version 4.0.2 using the functions *cor*.*test* and *prcomp* from the Base package respectively.

## Results

### In vitro 2-D cell migration exhibits conserved geometric features across cell line models

To investigate the differences in cell geometry during 2D-migration, we first defined quantitative morphological features which could be consistently extracted from *in vitro* imaging data. Due to its open-source availability, multi-operating system compatibility and user-friendly interface, the Fiji software was used in this study. Cell geometry features (CGFs) identified with the ‘Analyze Particles’ tool in Fiji were broadly categorized into size (Area, Perimeter, Width Height, Feret Diameter), shape (Circularity, Solidity, Roundness, Aspect Ratio) and position descriptors (X/Y co-ordinates, Feret Angle) as detailed in **Figure** Prior to examining trends during migration, we evaluated the statistical relationships between these CGFs using a migration dataset for high grade serous ovarian cancer (HGSC) cell lines^16^. The HGSC data comprised of six cell lines A4, OVMZ6, OVCA420, PEO14, OVCAR3, CAOV3 which were exposed to four distinct culture conditions during the *in vitro* wound healing assay *viz*., in the presence of 5% serum, serum starvation, exposure to chemotherapeutic drug paclitaxel (IC_50_ concentration as previously defined for individual cell lines^16^) and 0.1% DMSO as a drug-associated vehicle control. Quantification of wound closure over the course of the time-lapse experiment (16 hours) reiterated previous observations of efficacy in the following order A4> OVMZ6 > OVCA420 > PEO14 > OVCAR3 > CAOV3 (**Figure 2a**).

**Figure 1.**
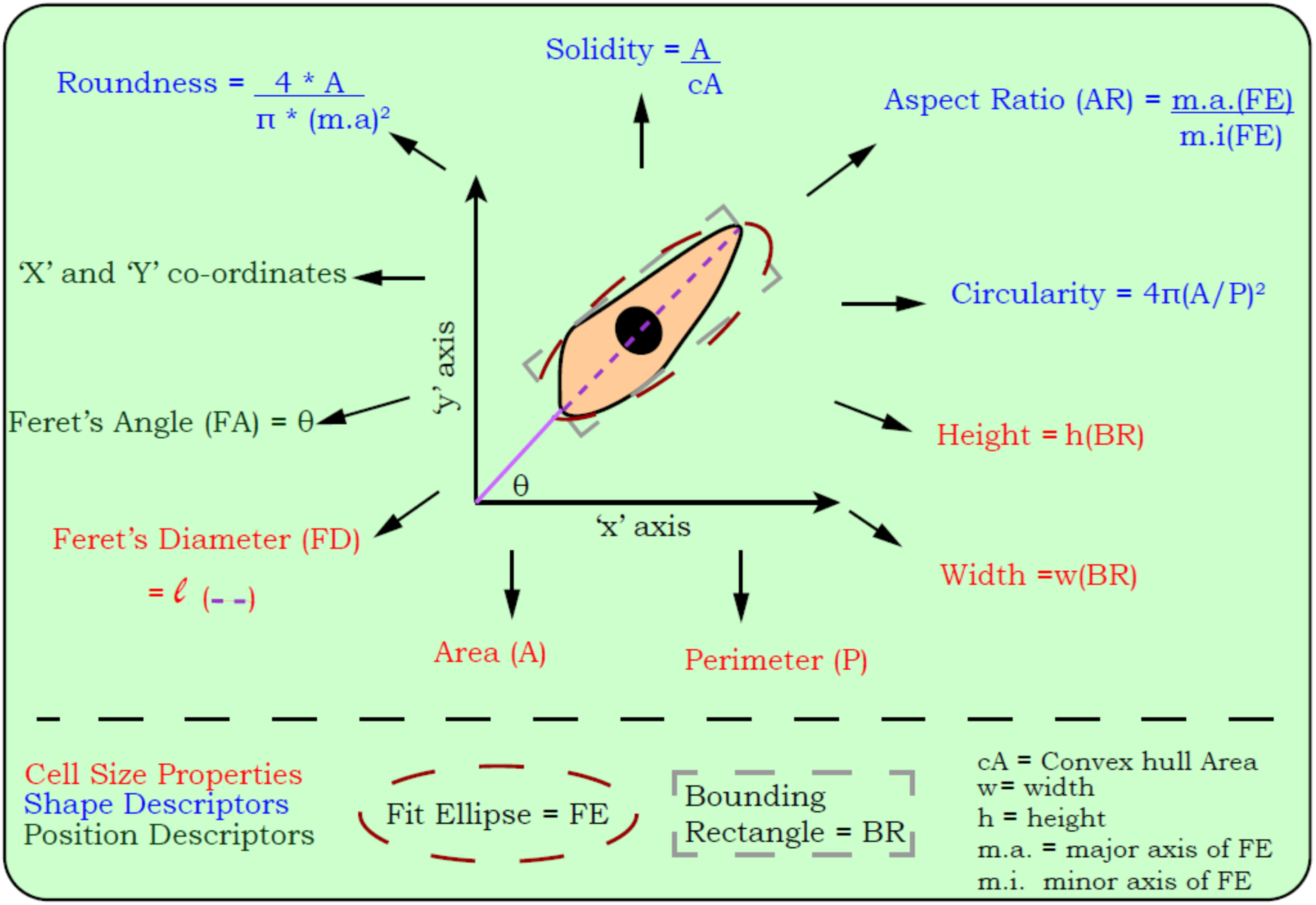
Cell Geometry Features. Schematic depiction of cell geometry features (CGFs) quantified for individual cells in migration datasets. CGFs were broadly classified into three categories *viz*., cell size, cell shape, and cell position. ‘Area’, ‘Perimeter’ and ‘Position Co-ordinates’ for each cell were directly quantified while other CGFs were derived by an artificial overlay of ‘fit ellipse’ and ‘bounding rectangle’ by the Fiji software. Formulae employed for quantifying CGFs by Fiji are outlined in the schematic.

**Figure 2.**
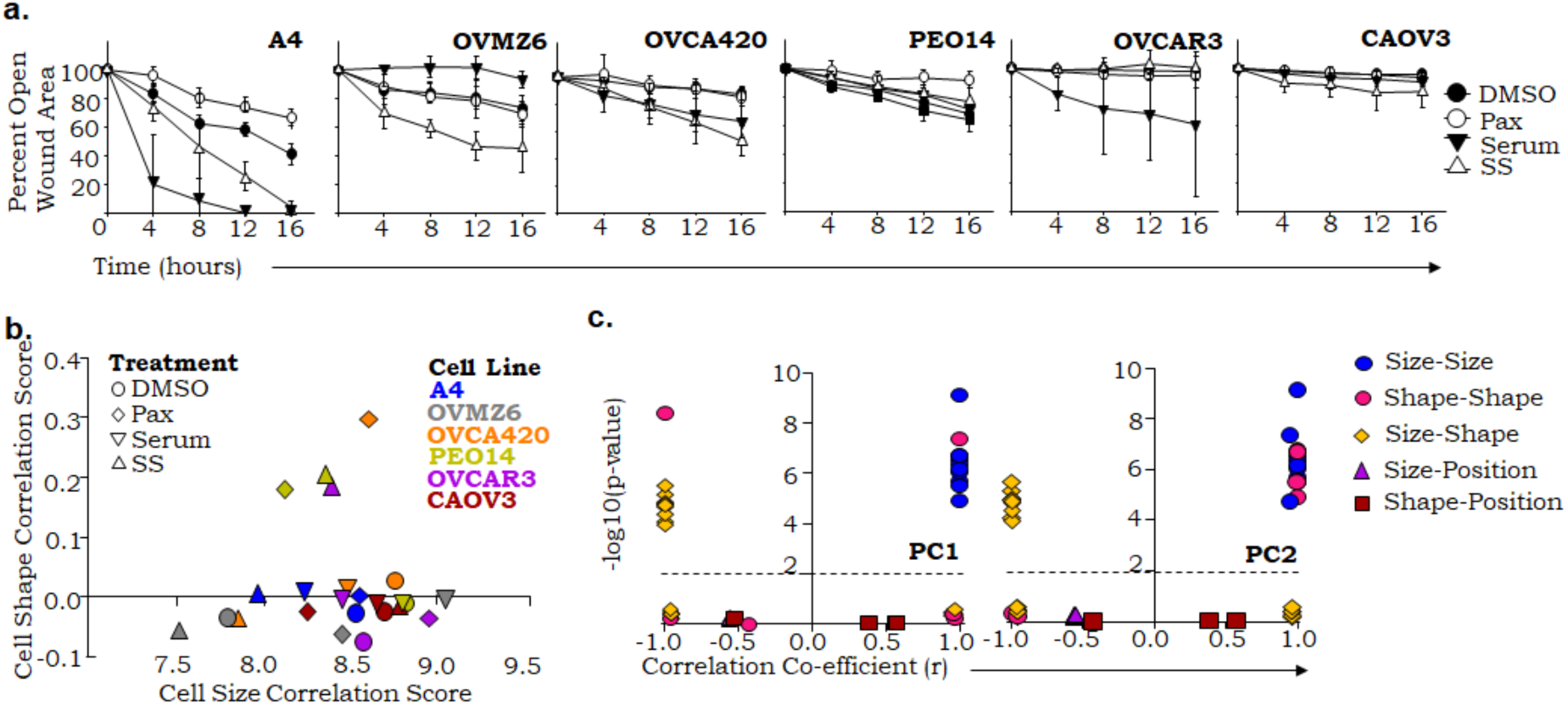
Conserved statistical relationships between cell geometry features are characteristic of 2D-cell migration. **a**. Percent open wound area quantified for a panel of high grade serous ovarian cancer (HGSC) cell lines in response to distinct assay conditions: DMSO, Paclitaxel (Pax), Serum, Serum Starved (SS). **b**. Cumulative correlation score-based (*detailed in Materials and Methods*) distribution of samples. **c**. Correlation coefficients computed for CGF-specific Eigen vectors for PC1 and PC2. Correlation coefficients were broadly calculated for five types of CGF pairs: (i) Size-size, (ii) shape-shape, (iii) size-shape, (iv) size-position and (v) shape-position. Except for panel (a), the data has not been segregated temporally and includes CGFs for individual cells from the HGSC migration.

We previously associated the phenotypic properties of each HGSC cell line with their migration capacity^16^; however, the dynamic changes in cell geometry were beyond the scope of the study. To obtain a preliminary understanding of their statistical inter-dependencies, pairwise correlation co-efficient between the ten CGFs were quantified without temporal segregation of data points. This revealed a significant positive correlation among cell size descriptors and independently, between the two cell shape descriptors ‘circularity’ and ‘solidity’. Further, these two CGF subsets (shape and size) negatively correlated with each other, but demonstrated no significant correlations with aspect ratio (AR), roundness and ferret angle (FA). While roundness and AR were negatively correlated, none of the CGFs correlated significantly with FA suggesting that CGF associations may be independent of cell orientation in a migrating monolayer (**Supplemental Figure 1**). To consolidate CGF correlative trends across multiple samples, we developed a cumulative correlation score (CCS) for cell size and shape descriptors separately (*Outlined in Materials and Methods*); FA was excluded from the analysis due to aforementioned observations. Distribution based on CCSs generated a tight cluster of samples with few outliers, despite the distinct phenotypic differences between cell lines (**Figure 2b**). To examine whether these distributions were affected over time during cell migration, we quantified the CCS across 30-minute intervals for each cell line-treatment pair and obtained comparable outcomes to what was seen in the earlier analysis (**Supplemental Figure 2**). These observations were conserved across the entire dataset, irrespective of the cell type or treatment, thus demonstrating the time- and position-independent nature of associations among the CGFs, which provided a robust framework for further analysis.

Next, we examined the contribution of individual CGFs to the variance and heterogeneity of cell migration. Principal component analysis (PCA) on temporally unsegregated CGF data reiterated the non-significant contribution of FA, hence cell orientation, to geometry (**Supplemental Figure 3a**). Principal components 1 and 2 (PC1 and PC2) captured approximately 65% of the total variance with the size descriptors, circularity, solidity and AR, roundness contributing to maximum variance in PC1 and PC2, respectively (**Supplemental Figures 3b, 3c**). Contribution of individual CGF pairs to PC axes was reminiscent of previous correlative trends wherein FA exhibited no quantitative significance (**Figure 2c**).

These observations resulted in the exclusion of FA from subsequent analysis and established the fundamental nature of CGF associations.

### Cell shape heterogeneity represents the plasticity of migratory cells

While we observed a consistent pattern of CGF associations, it was unclear whether these resulted from an averaging of heterogeneous sub-populations or the absence of phenotypic heterogeneity within cell lines. To examine the first hypothesis, individual cells across the HGSC migration dataset were assigned a motility status *viz*., non-migratory (NM) and migratory (M), based on the minimal distance covered by them during the experiment. The proportion of ‘NM’ and ‘M’ cells across the dataset exhibited cell line and treatment specific differences, with a majority of the cells being assigned the ‘NM’ phenotype (**Figure 3a**). Correlative trends quantified for ‘NM’ and ‘M’ cells were strikingly similar to those observed in the whole population data, thus affirming the absence of normalization effects (**Supplemental Figure 4, Supplemental Table 1**). Next, we tested whether the conserved CGF correlations emerged from the lack of morphological differences between the ‘NM’ and ‘M’ subsets. Examination of mean CGF values revealed that ‘M’ cells were larger in size and less circular than the ‘NM’ subset (**Figure 3b, Supplemental Figures 5 and 6**). While observations pertaining to circularity of ‘M’ cells were unsurprising, it was interesting to note the relatively small cell sizes of the ‘NM’ population.

**Figure 3.**
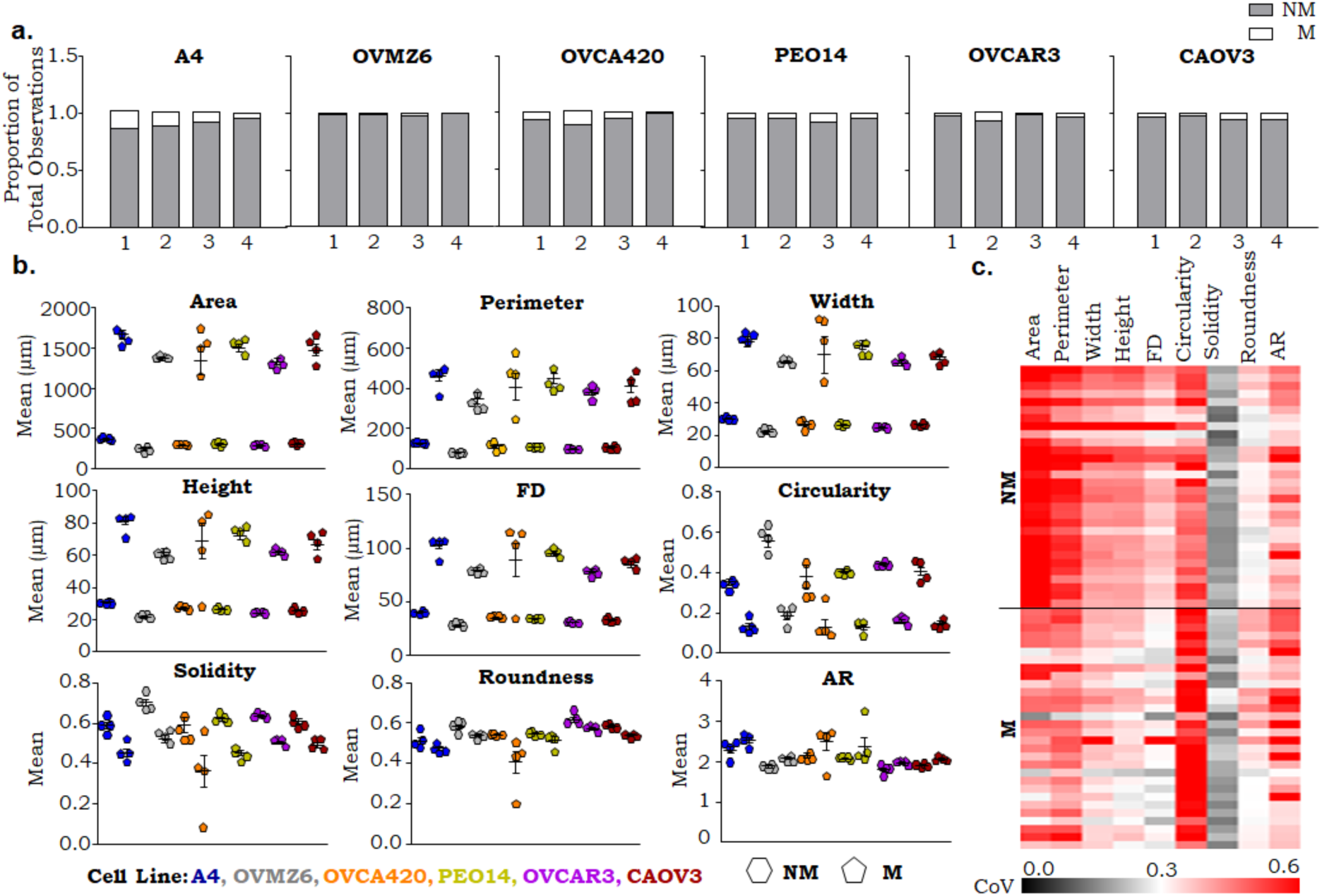
Consistent geometric trends identify heterogeneity in cell size for non-migrators and cell shape for migrators. **a**. Proportion of non-migratory (NM) and migratory (M) cells across the HGSC migration dataset. 1= DMSO, 2= Paclitaxel, 3= Serum, 4= SS. **b**. Mean distribution of CGFs between NM and M cells depicted for individual cell lines. Each data point depicts a specific treatment for individual cell lines. **c**. Heatmap depicting the co-efficient of variance for individual CGFs between NM and M cells. Data are not temporally segregated.

To further examine the heterogeneity of our cell systems, we evaluated the co-efficient of variance (CoV) for individual CGFs in the ‘NM’ and ‘M’ subsets. ‘NMs’ displayed a previously unappreciated heterogeneity in cell size which may be an outcome of differential proliferation and/or migratory-cell induced packing. On the other hand, the ‘M’ subset demonstrated greater heterogeneity of cell shape further reflecting its morphological plasticity (**Figure 3c, Supplemental Table 2**). The observed differences in CGF trends between ‘NM’ and ‘M’ cells thus, confirm the universal conservation of correlative trends and identify greater heterogeneity in functional subsets during 2-D cell migration.

### CGF features are conserved in an independent system of 2D monolayer migration

Robust statistical CGF trends prompted us to investigate whether similar observations could be extrapolated to an independent dataset. Literature search for articles with *in vitro* time-lapse migration data, provided a large cohort. Studies were selected based on the availability of good quality imaging data, processability with Fiji, the presence of appropriate controls and existence of relevant technical replicates. To examine CGF trends during *in vitro* wound healing, we extracted data from the Zaritsky *et al*. study wherein the wound healing efficacy of a mouse mammary adenocarinoma cell line (DA3) was examined in response to the growth factor HGF-SF. Differences in the cell system and, experimental setup permitted an unbiased assessment of CGF associations^25^. Data were not temporally segregated based on previous observations and analysed between control and HGF-SF treated samples (**Supplemental Figure 2**). Assignment of motility status identified a large population of ‘M’ cells in the control samples which were significantly enriched in response to HGF-SF (**Figure 4a**). A similar trend of large, non-circular ‘M’ cells as compared to the ‘NM’ subset was observed in both, control and HGF-SF treated samples (**Figure 4b**). Interestingly, increase in cell size for both, ‘NM’ and ‘M’ subsets along with reduced circularity for the non-migrators was evident following HGF-SF treatment. Changes in circularity can be attributed to an ‘NM’ to ‘M’ transition which explains the enhanced wound healing capacities in response to HGF-SF^25^. Quantifying CoV for DA3 cells revealed greater heterogeneity in ‘NM’ size as compared to the ‘M’ subset (**Figure 4c**). These observations can be explained by the existence of ‘M’ cells which have achieved homogeneity through extensive morphological changes as opposed to ‘NMs’ which are in the process of acquiring migratory properties. Thus, the Zaritsky, *et al*., dataset validated our CGF trends and provided an *in vitro* model for examining ‘NM’ to ‘M’ transition.

**Figure 4.**
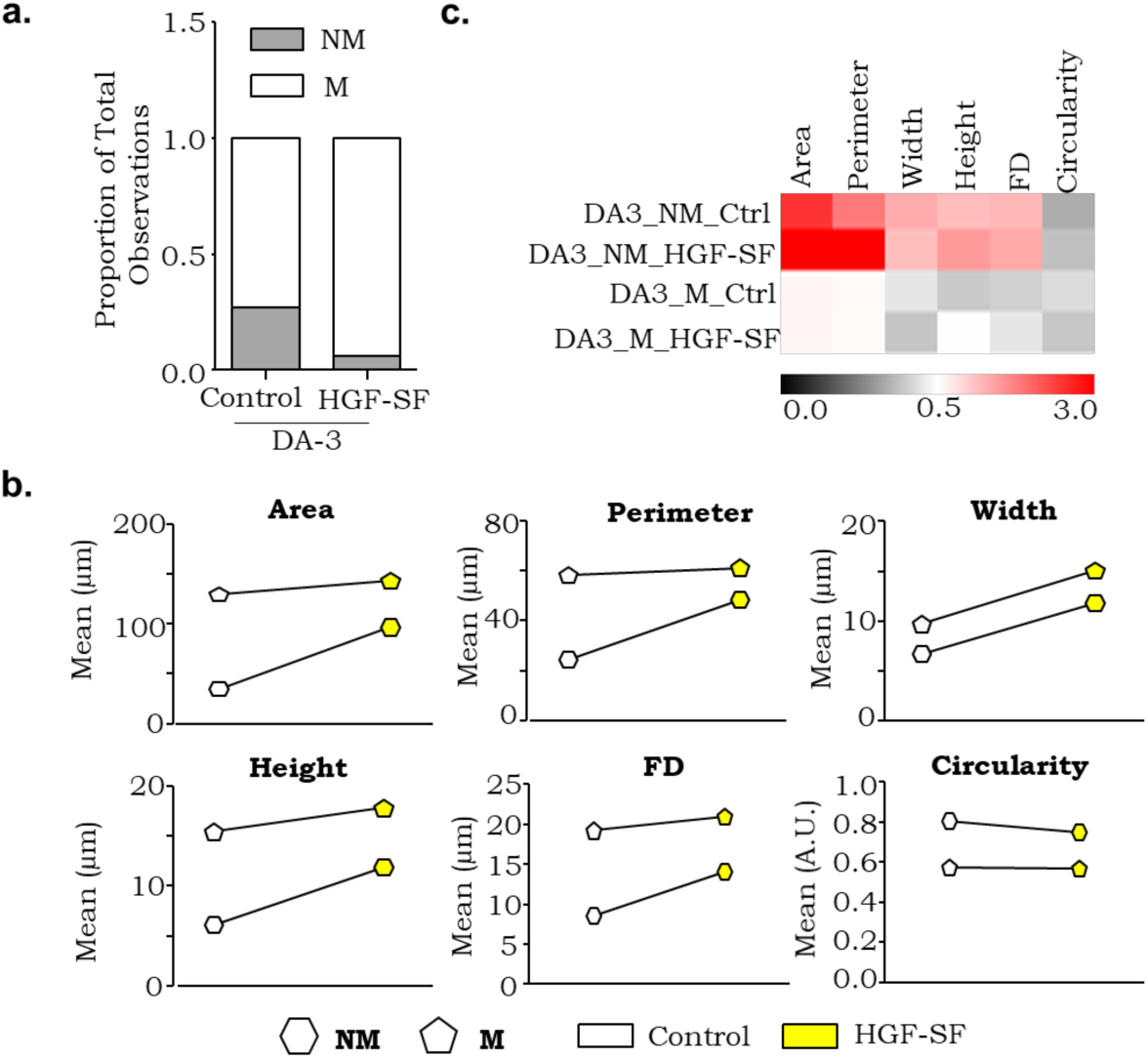
Mouse DA3 cells exhibit extensive non-migrator cell size variability. **a**. Proportion of non-migratory (NM) and migratory (M) cells in response to HGF-SF treatment. **b**.Mean distribution of CGFs between the NM and M cells. **c**. Heatmap depicting the trend for co-efficient of variance for individual CGFs between the NM and M cells. Data are not temporally segregated. Dataset represented: Zaritsky, et al.

### Sparsely seeded cells exhibit greater geometric plasticity in non-migratory as compared to migratory subset

We then investigated the utility of CGF trends in a dataset where directional migration, F-actin organization and focal adhesion maturation were examined in response to anisotropic substrate alignment^26^. Due to differences in image quality, we only extracted data for individually migrating T47D and KPC-derived cells. Annotation of migratory subsets revealed a higher proportion of ‘M’ cells in KPC as compared to T47D (**Supplementary Figure 7a**). Unlike previous observations, CGF trends demonstrated a cell line specific trend with T47D and KPC exhibiting larger cell size in ‘M’ and ‘NM’ subsets, respectively (**Supplementary Figure 7b)**. Both systems demonstrated heterogeneity in size with relatively homogenous cell shape features irrespective of the migratory subset analysed (**Supplementary Figure 7c)**. While these observations may be an outcome of the experimental setup, they could also result from differences in cell packing which is non-existent in sparsely seeded, individually migrating cells.

### 2D packing contributes to CGF variability in a system of confluent monolayer migration

To examine the effects of 2D-packing on cell geometry, we used a dataset where the biophysical dynamics of collective migration were studied on confined PolyDiMethylSiloxane (PDMS) islands ^24^. Data were extracted for MDCK cells subject to three experimental setups *viz*., cytochalasin ‘D’ treatment (CytoD), high cell density (High) and low cell density (Low). While a higher proportion of ‘M’ cells was observed in all samples, their frequency increased in the following order CytoD < High < Low (**Figure 5a**). These observations could be explained by the inhibitory effects of CytoD on cytoskeletal re-organization and effects of cell density on 2D-packing and -motility. Unlike previous data, CGF trends were reversed for ‘NM’ and ‘M’ subsets. Large, less circular ‘NMs’ were observed for both ‘High’ and ‘Low’ experimental groups while ‘CytoD’ did not significantly affect either migratory subset (**Figure 5b**). Differences in CGF trends can be explained by the extent of ‘NM’ spreading, potentially facilitated by collagen coated growth surfaces which were absent in the HGSC and DA3 datasets. CoV analysis identified more heterogeneity in ‘NMs’ versus ‘Ms’ with ‘CytoD’ samples exhibiting highest variability in NM size. While cell size variability can be explained as an outcome of CytoD treatment, the greater homogeneity of Ms may be attributed to multiple factors some of which include cell system bias, collagen/ECM-driven uniformity in cell shape and restricted migratory spaces due to NM spreading resulting in limited morphological conformations (**Figure 5c**). Due to the use of a cell monolayer in this data, the CGF trends would principally depend on the extent of cell packing which are influenced by cell division/cell death events, and the frequency of NM-M transitions. These observations highlight the influence of spatial constraints on, driving CGF trends and differentially affecting the migratory subsets.

**Figure 5.**
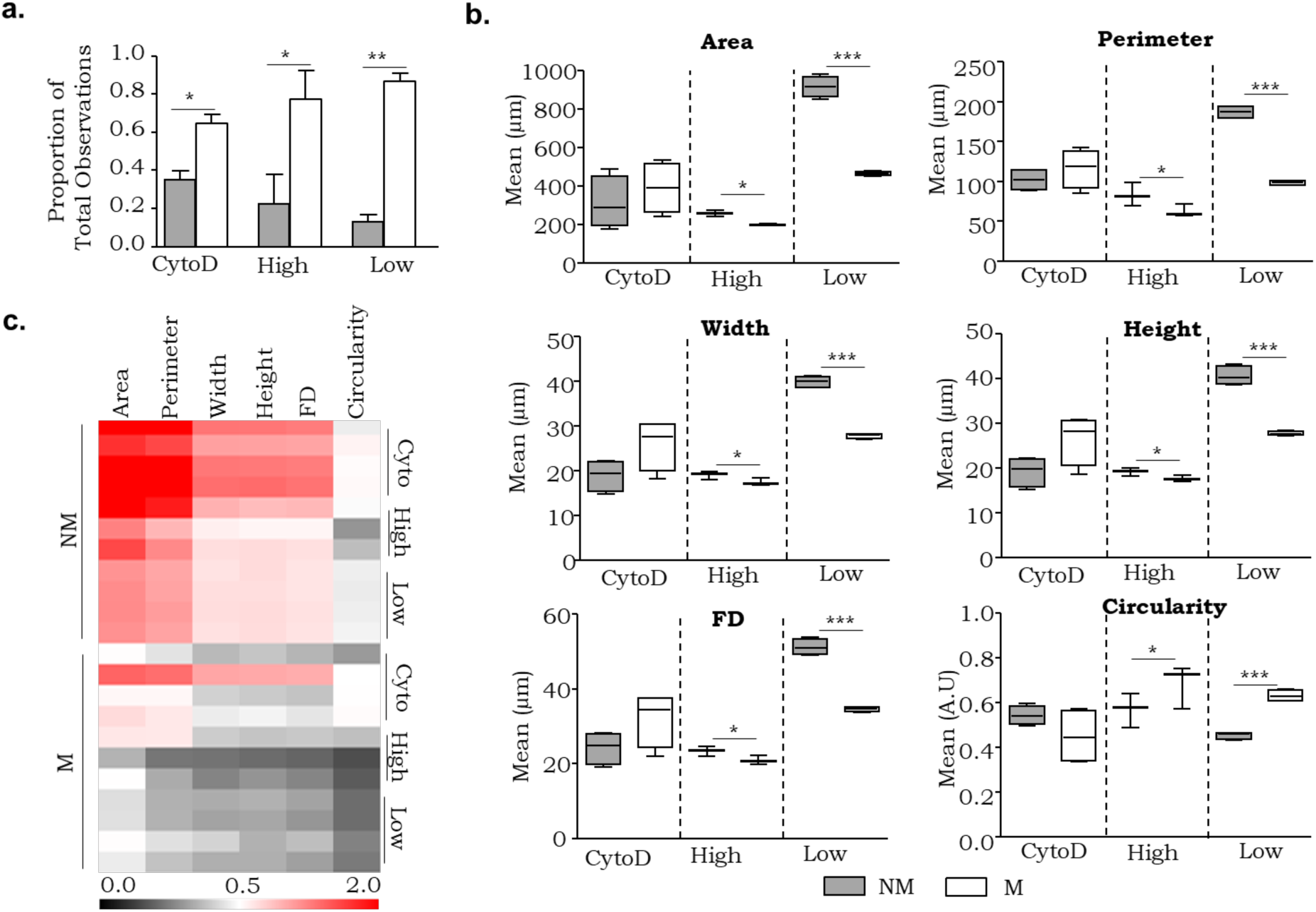
Migration associated CGF trends in a confluent monolayer. **a**. Proportion of non-migratory (NM; grey) and migratory (M; white) cells in response to Cytochalasin D (CytoD) treatment, high density (High) and low density seeding (Low) on micropattern wells. **b**. Mean distribution of CGFs between NM and M cells. **c**. Heatmap depicting the co-efficient of variance for individual CGFs between the NM and M cells. Data are not temporally segregated. Dataset represented: Saraswathibhatla, et al.

## Discussion

Decades of research on cell migration and recent developments with *in vitro* assays have been fruitful in understanding several biological processes and for evaluating therapeutic strategies. Due to their widespread utility, analytical pipelines that augment migration read outs have received plenty of attention. In the present study, we have designed a quantitative approach based entirely on open source algorithms for label-free time lapse migration data and applied it for the resolution of functional heterogeneity during migration. Cell geometry was the key parameter evaluated in this analysis and its associated features exhibited consistent trends across experimental systems (**Supplemental Figures 2 and 8**). Our pipeline allows the evaluation of migratory subsets in response to extrinsic cues, wound induction, 2D-packing and provides a foundation to understand migratory transitions.

Our primary dataset for the development of this approach was a panel of HGSC cell lines, where we had previously identified distinct migratory modalities at the population level^27^. Our previous approach had precluded the heterogeneous behaviour of individual cells and exclusively focused on cells that undertook quantifiable displacement during migration. Incorporation of CGFs at a single cell resolution demonstrated the fundamental geometric differences between migratory and non-migratory subsets across phenotypically diverse cell systems including independent datasets; importantly, these trends were unaltered by exogenous cues^25,27,28^. We also observed an unprecedented heterogeneity in the non-migratory subset which may result from one or more of the following reasons: cells existing in different, (i) states of proliferation^29^, (ii) functional states while transitioning between migratory subsets^30^, or (iii) a greater plasticity of cells with non-specialized functions^31^. The ability of our pipeline to discern proportion of NM to M cells (NM/M) describes the functional composition of cell systems and can prove useful in pharmacological screens. Our observations suggest that the pre-existence of specific migratory subsets and dynamics of their interconversions may be crucial for migration efficiency.

Predictive models designed around tissue- and assay-specific variables have suggested the role of cell geometry during *in vitro* migration^29^, metastatic invasion^32^, and cell polarization for morphogenesis^33^. Experimental evidence linking cell geometry with molecular compartmentalization^34^, intercellular crosstalk^35^, surface topographies^36^ and mechano-reciprocity^35^ have further indicated the utility of this parameter in distinguishing functionally distinct cells. Previous reports on colon and breast cancer have also utilized CGFs to predict the chemo-responsiveness^37^ and invasive potential^38^, respectively, in cell line models. Unlike published studies, our pipeline can be applied not only to scratch assay but also data from confluent monolayers and individual cells. The Saraswathibhatla *et al*., and Zaritsky *et al*., datasets were two such distinct studies which upon CGF analysis allowed us to speculate the role of 2D-packing on migratory heterogeneity^24,25^. While we cannot ignore the cell and experimental system specific effects, these trends between migratory subsets do recapitulate jamming-unjamming transitions which are reported during tissue development and disease progression^8,39,40^. While not entirely evident from our analysis, previous reports have provided evidence of cell state transitions in response to spatial packing^41,42^.

Our analytical pipeline relies on open source free software which can be used across recent computing systems, irrespective of their processing capacity. The application of our approach to label-free systems reduces the dependence on sophisticated labelling or microscopy techniques to capture cell migration. However, due to its reliance on phase differences for distinguishing cells, the quality of input data and presence of debris in the field of view can affect the analysis. We trust that the inclusion of fluorescence microscopy approaches will enhance the sensitivity of our approach and may further resolve CGF trends. We are aware of several image analysis pipelines developed specifically to study cell migration, with each approach presenting a distinct set of advantages^13^. We believe that our pipeline builds on existing quantitative measures and provides a simple analytical platform to dissect the functional heterogeneity during 2D migration. We acknowledge the speculative nature of our conclusions and the need to experimentally validate transitions between NM and M subsets, as well as spatio-temporal specific patterns of NM and M subpopulations. While we initially intended to examine the temporal and spatial heterogeneity influencing migration dynamics, we were unable to capture this information with our current approach, thereby limiting our capacity to investigate the degree of influence of neighbouring cells on migratory behaviour.

Overall, we have developed an image analysis pipeline and demonstrated its utility across experimental systems for identifying distinct migratory subsets. While future development will add to its value, we believe that in its current form our analytical approach will greatly benefit biologists interested in the comprehending the process of cell migration.

## Acknowledgements

The authors thank all members of Jolly lab for discussion and input for this manuscript. M.K.J. was supported by Ramanujan Fellowship awarded by Science and Engineering Research Board (SERB), Department of Science & Technology, Govt. of India (SB/S2/RJN-049/2018).

## Author Contributions

S.S.V. and M.K.J. conceived research. M.K.J. (computational) and S.A.B. (experimental) supervised research. S.S.V. and K.H. performed research and analysed the data. All authors contributed to manuscript writing and have read and agree to this version of the manuscript.

## Data and Code Availability

Raw data for the HGSC migration dataset is available upon request from S.A.B. Raw data from other datasets used in this study was extracted from the data submitted along with the original publications. Raw data for CGFs is available on the drive link: https://drive.google.com/drive/folders/1gmxfFJyNkyuxmWW3rO623qbgiC8251Vd?usp=sharing. Codes used for structuring, analysing, and visualising the data are available on GitHub at https://github.com/askhari139/CGF.

