## Supplementary Information for "Cell Geometry Distinguishes Migration-Associated Heterogeneity in Two-Dimensional Systems"

**Supplemental Information**

**Supplemental Figures**

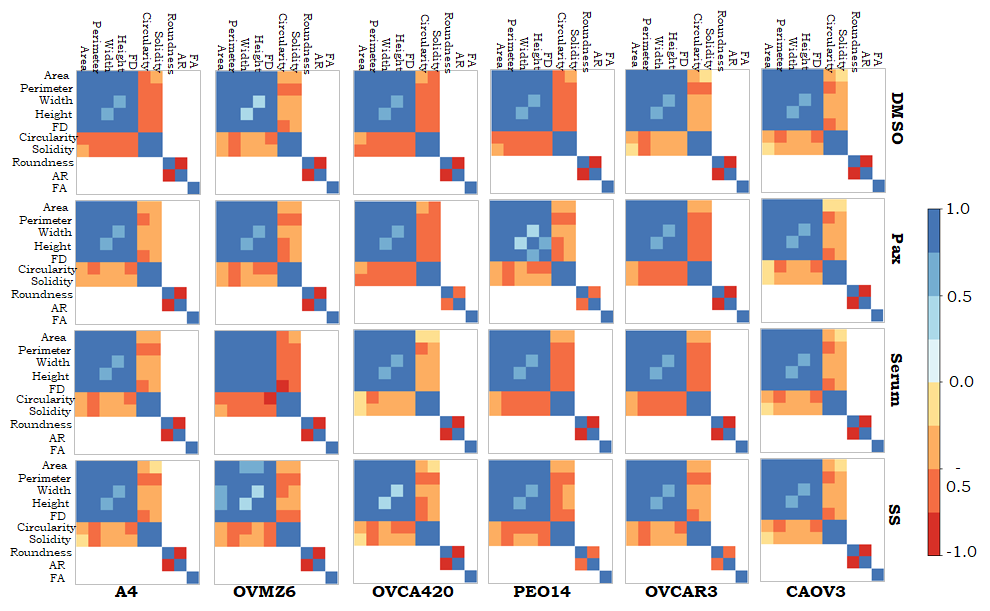

**Supplemental Figure 1. Correlative trends between CGFs are conserved across a panel of HGSC cell lines.** Heat-maps depicting significantly correlated CGFs (*P*<0.05) across the HGSC migration dataset. Non-significant correlations are represented in white. Data represented in this figure is not temporally segregated. FD: Feret Diameter, AR: Aspect Ratio, FA: Feret Angle, DMSO: Dimethyl Sulfoxide, Pax: Paclitaxel, SS: Serum starved.

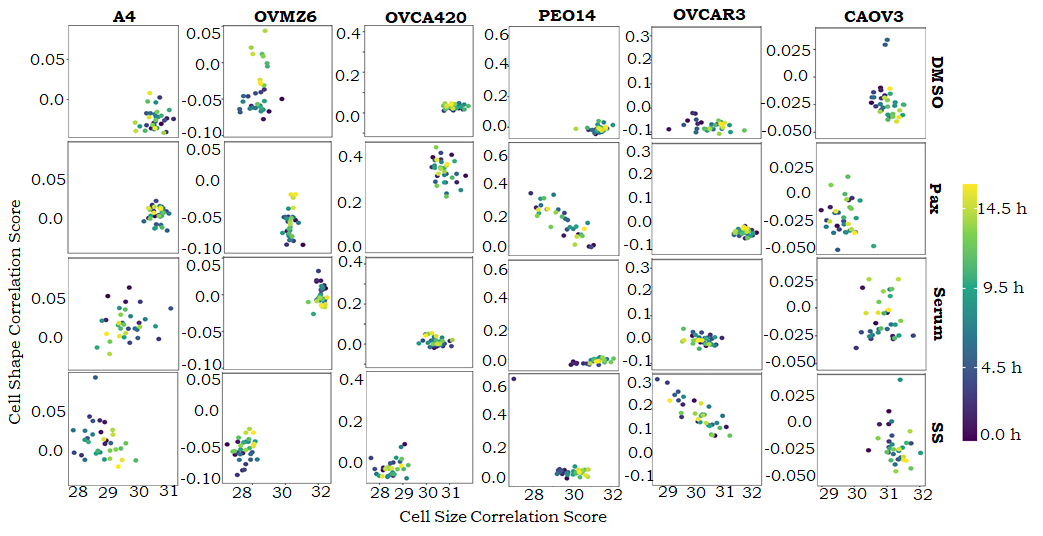

**Supplemental Figure 2. Correlation scores are temporally conserved across the HGSC cell line panel.** Scatter plots depicting the correlation score-based distribution of samples. Individual dot depicts data from a distinct time point. Time scale is represented as a gradient from 0 – 16 hours following wound induction. DMSO: Dimethyl Sulfoxide, Pax: Paclitaxel, SS: Serum starved.

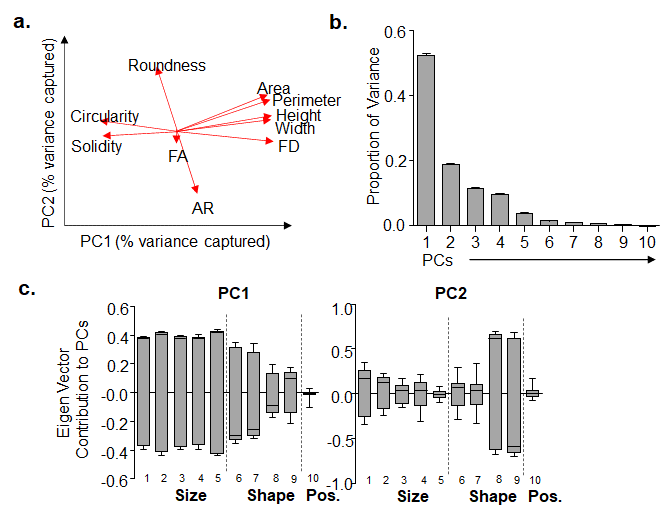

**Supplemental Figure 3. CGF Variance. a.** Schematic depicts the contribution of individual CGFs to percent variance captured across principal components 1 (PC1), and 2 (PC2). **b.** Proportion of variance captured by individual PCs (PC1-PC10). **c.** Eigen vector contributions of individual CGFs to the variance captured by PC1 and PC2. Eigen vectors derived for each CGF are represented by **Size-** 1: Area, 2: Perimeter, 3: Width, 4: Height, 5: Feret Diameter (FD); S**hape-** 6: Circularity, 7: Solidity, 8: Roundness, 9: Aspect Ratio (AR); **Position**- 10: Feret Angle (FA). Data represented in this figure is not temporally segregated and each graph is representative of data obtained from the entire HGSC migration dataset.

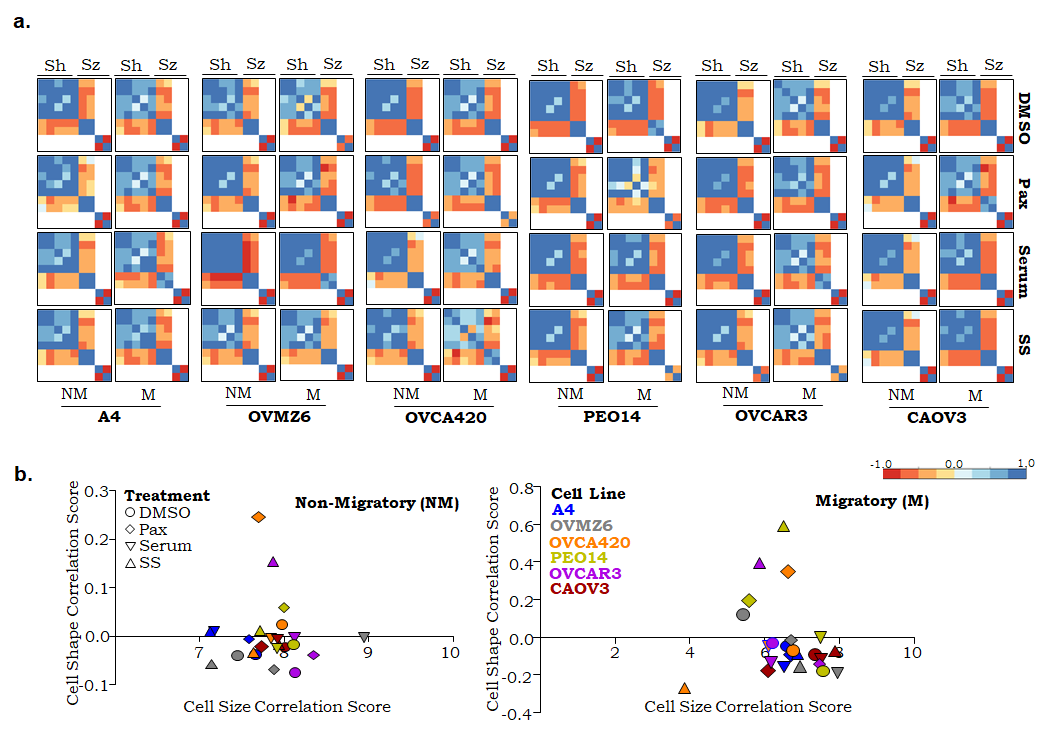

**Supplemental Figure 4. Correlative trends for CGFs are conserved across migratory and non-migratory sub-populations. a. H**eat-maps depicting significantly correlated CGFs (*P*<0.05) across the HGSC migration dataset. Non-significant correlations are represented in white. Data represented in this figure has not been temporally segregated. Sh: Shape Descriptors, Sz: Size Descriptors, NM: Non-migratory, M: Migratory. **b.** Correlation score-based distribution of samples. DMSO: Dimethyl Sulfoxide, Pax: Paclitaxel, SS: Serum starved.

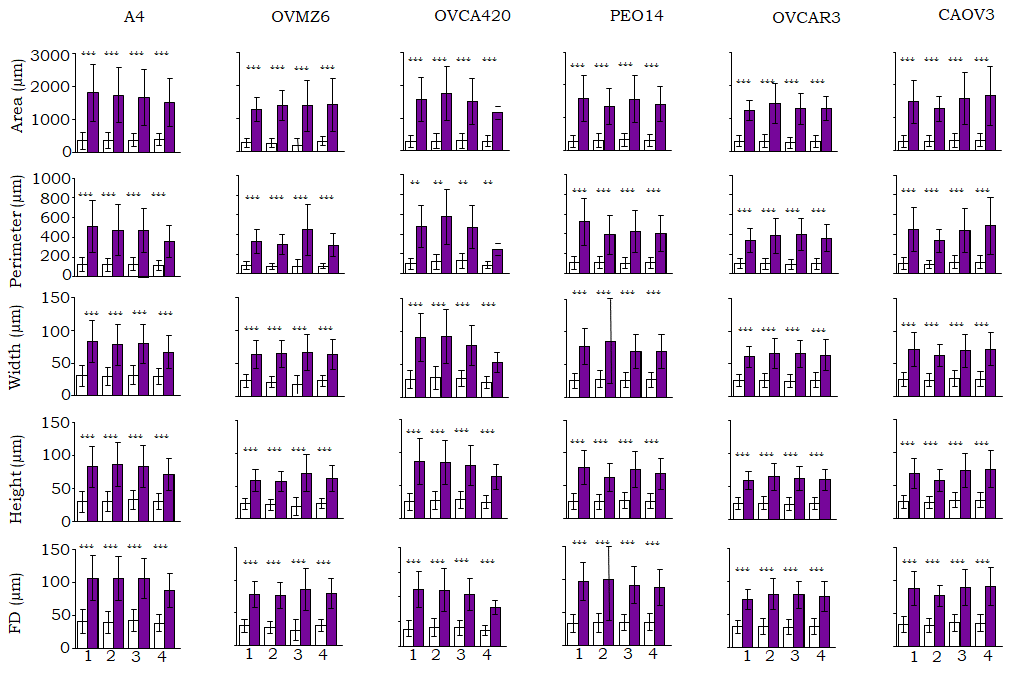

**Supplemental Figure 5. Migratory cells are larger than non-migratory cells.** Quantitative comparison of cell size CGFs between non-migratory (white) and migratory (purple) cells in the HGSC migration dataset. Cells were subjected to 4 different assay conditions, 1: DMSO, 2: Paclitaxel, 3: Serum, 4: Serum Starved. Data depicted in the figure is not temporally segregated. FD: Feret Diameter. All data are depicted as mean ± SEM. *P < 0.05, **P < 0.01, ***P < 0.001.

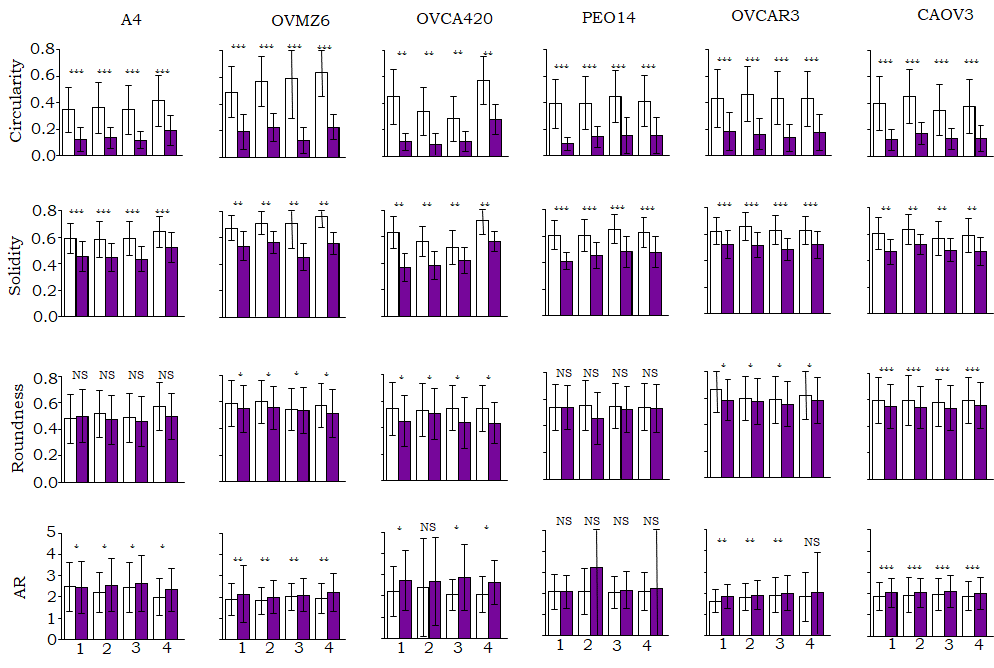

**Supplemental Figure 6.** **Migratory cells adopt non-circular shapes as opposed to non-migrators.** Quantitative comparison of cell shape CGFs between non-migratory (white) and migratory (purple) cells in the HGSC migration dataset. Cells were subjected to 5 different assay conditions, 1: DMSO, 2: Paclitaxel, 3: Serum, 4: Serum Starved. Data depicted in the figure is not temporally segregated. AR: Aspect Ratio. All data are depicted as mean ± SEM. *P < 0.05, **P < 0.01, ***P < 0.001.

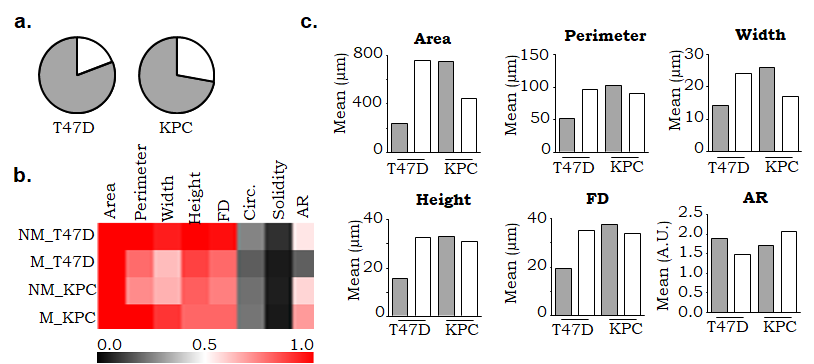

**Supplemental Figure 7. CGF trends in individually migrating cancer cells. a.** Proportion of non-migratory (NM; grey) and migratory (M; white) cells in the Ray *et al.*, dataset. **b.** Heatmap depicting the co-efficient of variance for cell size and shape descriptors in NM and M cells. **c.** Quantitative comparison of CGFs between NM (grey) and M (white) cells. Dataset represented: Ray, *et al*.

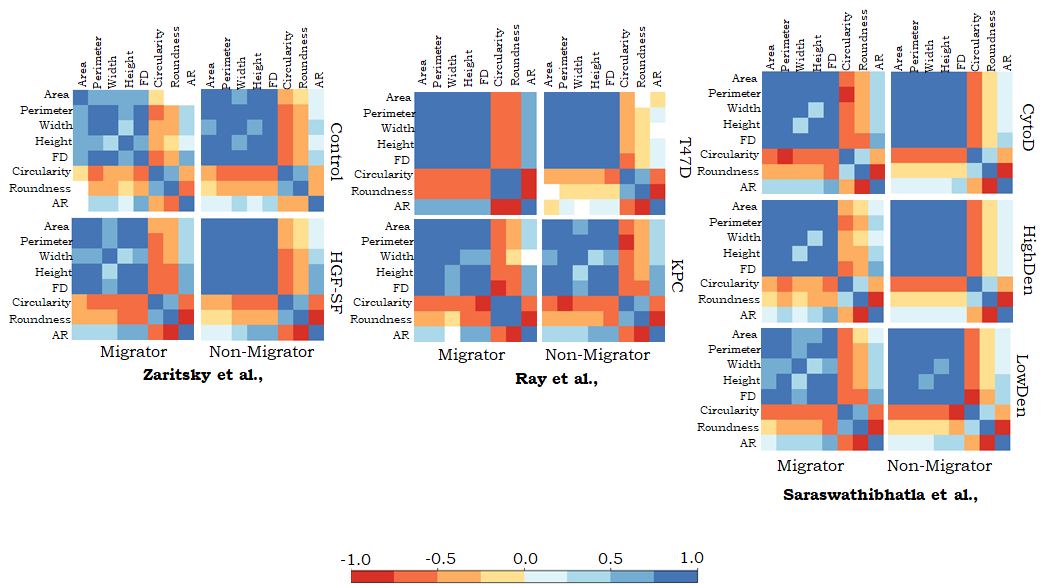

**Supplemental Figure 8. CGF correlative trends across independent datasets.** Heat-maps depicting significantly correlated CGFs (*P*<0.05) across the Zaritsky *et al.*, Ray *et al.*, and Saraswathibhatla *et al.*, datasets. Non-significant correlations are represented in white. Data represented in this figure is not temporally segregated. FD: Feret Diameter, AR: Aspect Ratio. Solidity was excluded from the analysis due to its low variance as depicted in **Figure 3c.**

**Supplemental Tables**

*Supplemental Table 1: Raw values for correlation co-efficient computed for all CGFs in whole-population (WP), non-migratory (NM) and migratory (M) subsets of the HGSC migration dataset. Data includes differences in correlation coefficient computed for (WP-NM), (WP-M) and (NM-M).*

*Supplemental Table 2: Coefficient of Variance (CoV) computed for non-migratory (NM) and migratory (M) cells in the HGSC migration dataset. Data includes a ratio for CoV computed for (NM/M).*
